# Diversity of fate outcomes in cell pairs under lateral inhibition

**DOI:** 10.1101/032078

**Authors:** Nara Guisoni, Rosa Martinez-Corral, Jordi Garcia Ojalvo, Joaquín de Navascués

## Abstract

Cell fate determination by lateral inhibition via Notch/Delta signalling has been extensively studied. Most formalised models consider Notch/Delta interactions in fields of cells, with parameters that typically lead to symmetry breaking of signalling states between neighbouring cells, commonly resulting in salt-and-pepper fate patterns. Here we consider the case of signalling between isolated cell pairs, and find that the bifurcation properties of a standard mathematical model of lateral inhibition can lead to stable symmetric signalling states. We apply this model to the adult intestinal stem cell (ISC) of *Drosophila*, whose fate is stochastic but dependent on the Notch/Delta pathway. We observe a correlation between signalling state in cell pairs and their contact area. We interpret this behaviour in terms of the properties of our model in the presence of population variability in signalling thresholds. Our results suggest that the dynamics of Notch/Delta signalling can contribute to explain stochasticity in stem cell fate decisions, and that the standard model for lateral inhibition can account for a wider range of developmental outcomes than previously considered.

**Summary statement:** Notch/Delta-mediated lateral inhibition in cell pairs can result in symmetric signalling depending on the activation threshold, which can modulate cell-fate decisions depending on contact area.

## Introduction

The Notch/Delta signalling pathway is one of the main regulators of cellular differentiation during development and adult tissue maintenance (reviewed in (Artavanis-Tsakonas et al., 1999; Ehebauer et al., 2006; Koch et al., 2013). It often drives mutually inhibitory interactions between cells, acting as a gate for differentiation. This mode of action has been termed lateral inhibition, and has been the object of experimental study as well as mathematical formalisation for decades (see, for instance, Othmer and Scriven, 1971; Collier et al., 1996; Sprinzak et al., 2011; Petrovic et al., 2014). Quantitative models of lateral inhibition usually involve a field of cells expressing initially similar amounts of the receptor Notch and its membrane-bound ligand Delta. Delta trans-activates Notch in neighbouring cells and Notch, once activated, reduces in turn the ability of the cell to signal through Delta, leading to a state of mutual repression. This symmetry (and cell fate equivalence) is eventually broken by enforced biases and/or stochastic variation in Notch/Delta levels (Collier et al., 1996; Plahte, 2001; reviewed in Simpson, 2001) resulting in extended finegrained spacing patterns (Othmer and Scriven, 1971; Collier et al., 1996; see also Shaya and Sprinzak, 2011) that have been experimentally characterized in depth in real developmental systems (reviewed in Greenwald, 1998; Arias and Stewart, 2002). In contrast, little attention has been paid so far to the effect of lateral inhibition in isolated cell pairs, beyond the trivial expectation that symmetry breaking will eventually take place, leading to cells taking opposing fates (see for instance, Collier et al., 1996; Rouault and Hakim, 2012). However there has been no formal investigation of whether alternative steady states are possible, perhaps due to the lack of an experimental model to relate it to.

The cellular homeostasis of the adult *Drosophila* midgut can provide this experimental scenario, as in this tissue Notch/Delta signalling occurs mostly in isolated pairs of cells (Ohlstein and Spradling, 2006; de Navascués et al., 2012; Goulas et al., 2012) (Fig.1A). The fly’s intestinal lining is maintained by intestinal stem cells (ISCs), which divide to both self-renew and provide committed progenitors. Progenitors specialise in producing either nutrient-absorbing enterocytes or secretory enteroendocrine cells (Micchelli and Perrimon, 2006; Ohlstein and Spradling, 2006; Zeng and Hou, 2015; Guo and Ohlstein, 2015). The precursors of enterocytes, called enteroblasts (EBs) are frequently found forming pairs with ISCs (Fig. 1A,B). These pairs are thought to result from an earlier division of an ISC and subsequent fate allocation by Notch signalling, before a new division or terminal differentiation event takes place (Goulas et al., 2012; de Navascués et al., 2012). Importantly, ISC divisions in the enterocyte lineage result in either asymmetric fate (one ISC and an EB), or symmetric self-renewal (two ISCs) or differentiation (two EBs), which globally result in balanced, homeostatic proportions (de Navascués et al., 2012) (Fig. 1C-E). This mode of tissue maintenance, whereby the balance between stem cell self-renewal and differentiation is achieved at the population level rather than within every stem cell lineage, is termed neutral competition (Klein and Simons, 2011) and is found in a growing number of self-renewing adult tissues (Simons and Clevers, 2011). While no molecular mechanism has been fully elucidated so far for any case of neutral competition, in the fly gut it has been proposed to arise from lateral inhibition mediated by Notch/Delta (de Navascués et al., 2012), a pathway known to define the fate of the ISC offspring (Ohlstein and Spradling, 2006; Micchelli and Perrimon, 2006; Bardin et al., 2010).

**Figure 1.**
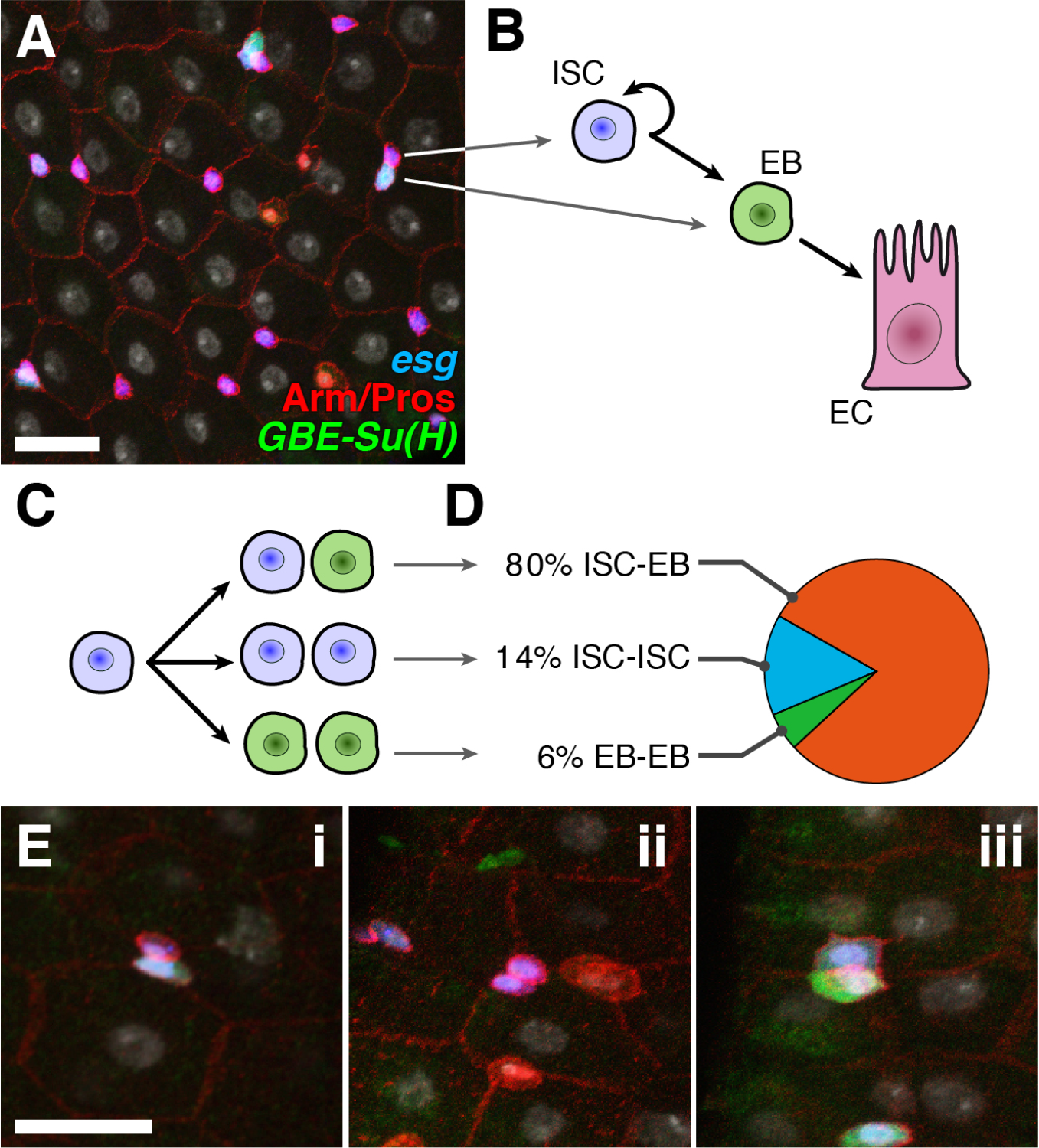
Tissue maintenance in the Drosophila adult midgut. Scale bars: 20*µ*m. A. Confocal micrograph showing the cell types present in the midgut epithelium. ISCs are esg-GFP+ (blue) and EBs are esg-GFP+ and GBE-Su(H)-lacZ+ (green). The two differentiated cells, enteroendocrine cells and enterocytes, are recognisable by Prospero (pros) expression and having large, polyploid nuclei (Hoechst, grey), respectively. B-C. ISCs self renew and produce EBs (which will terminally differentiate without further division) (B), by dividing either asymmetrically (one ISC and an EB), or symmetrically into two ISCs or two EBs (C). D. Distribution of cell fates for nests containing two undifferentiated cells (N=508). ISC-ISC pairs (blue, N=74), EB-EB pairs (green, N=28) and ISC-EB pairs (orange, N=406). E. Confocal micrographs showing examples of cell pair fate profiles: asymmetric (i), symmetric Notch negative (two ISCs, ii) and symmetric Notch positive (two EBs, iii).

Here we explore the capacity of a standard model of lateral inhibition acting in pairs of interacting cells to result in steady states with different signalling states (either symmetric or asymmetric) coexisting in the tissue. We find that this is indeed possible, provided there is population-wide variation of signalling thresholds. Next, we turn to the *Drosophila* midgut and find that the tissue displays high variability of contact area between pairs of ISC/EB cells, which can be associated to an effective heterogeneity in signalling thresholds between pairs of cells. When contrasting this variability with the distribution of fate combinations in pairs of ISC/EB cells, we find a correlation between contact area of specific cell pairs and their fate profile. Moreover our model is able to reproduce the distribution of fate outcomes given the contact area distribution.

Our results expand the repertoire of possible outputs of a system governed by lateral inhibition, and connect this mode of signalling with a mode of stem-cell based tissue maintenance (neutral competition) that is highly relevant in adult tissue homeostasis and tumourigenesis (Simons and Clevers, 2011; Vermeulen et al., 2013; Baker et al., 2014), and whose molecular regulation is poorly understood.

## Materials and Methods

### The model: lateral inhibition mediated by Notch-Delta interaction

We consider that the rate of Notch activation in a cell is an increasing function of Delta concentration on its neighbour (signalling), and that the rate of Delta expression is a decreasing function of the level of activated Notch in the same cell (inhibition). We represent these interactions by means of a standard mathematical model of Notch/Delta signalling (Collier et al., 1996) between pairs of cells, which is given by:

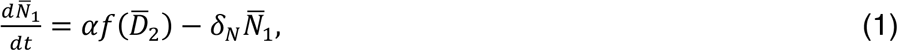

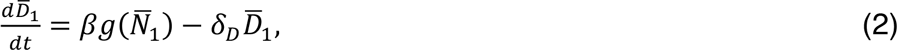

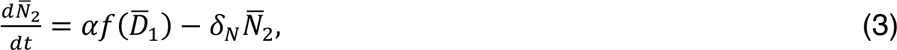

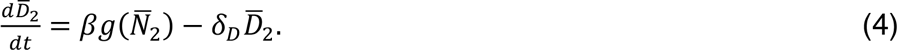

Here 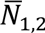 represent the levels of Notch activity in cells 1 and 2, and 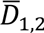 are the concentrations of Delta in each cell. α and β are the maximal production rates of Notch and Delta, respectively, whereas *δ_N_* and *δ_D_* are their corresponding degradation rates. The production terms for Notch (*f*) and Delta *(g)* are given by the Hill functions

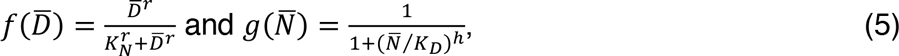

where the first function represents the signalling effect of Delta on the neighbouring cell, and the second corresponds to the inhibition of Delta expression by activated Notch in the same cell. *K_N_* is the threshold of Notch activation by neighbouring Delta, *K_D_* is the threshold of Delta inhibition by Notch in the same cell, and the coefficients *r* and *h* represent the cooperative character of the two aforementioned processes. Similarly to Collier et al, (1996) we rewrite Eqns 1-4 in dimensionless form as:

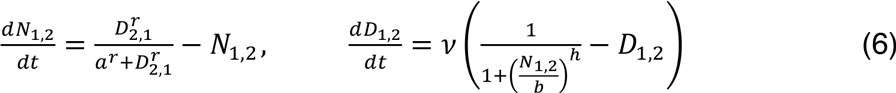

The parameter *ν* is the ratio between the degradation rates of Delta and Notch, *δ_D_* /*δ_N_*. *a* and *b* are the dimensionless thresholds for Notch activation by Delta in the neighboring cell, and Delta inhibition by Notch in the same cell, respectively,

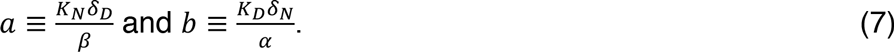

*a* and *b* are referred to as the activation and inhibition thresholds, and their values set the location of the half-maximal points of the Hill functions in Eqn 6.

### Steady states and cell fate identification

The system of equations from Eqn 6 has a homogeneous steady state in which Notch and Delta have the same values in the two cells:

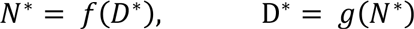

This state corresponds to a situation in which both cells in the pair have the same fate. The stability boundary of this homogeneous steady state can be calculated using standard methods (Collier et al., 1996), and is represented by a dotted grey line in Fig. 2A. Above this line the homogeneous state is stable. Below it, a heterogeneous stable steady state appears in which the values of Notch and Delta are different between the two cells:

**Figure 2.**
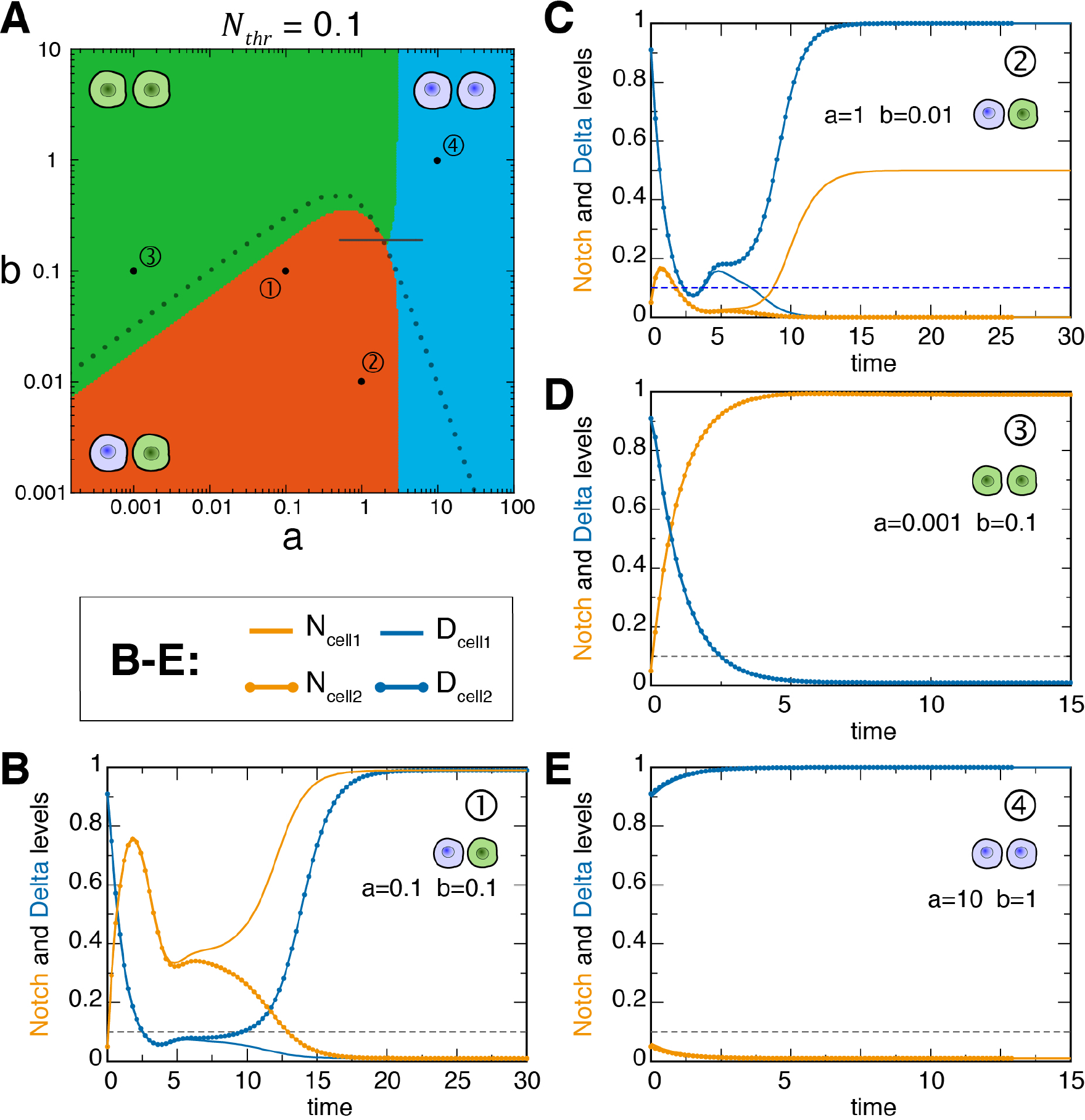
Parameter space and dynamic behaviour of the model. A. Stable solutions of the system classified according to their resulting signalling state. Green stands for symmetric positive fates (EB-EB pairs), blue represents symmetric negative fates (ISC-ISC pairs), and orange denotes asymmetric fates (ISC-EB pairs). The threshold in Notch level for EB identification is taken to be equal to 0.1 (see the text for more details). The short, horizontal line indicates 95% of “a” values used to generate a theoretical distribution of cell fates for b=0.19 (see main text). Dotted line, boundary of stability for steady states with identical cells; these ‘homogeneous’ solutions are stable above the line. B-E. Time evolution (in arbitrary units) of Notch and Delta activity in pairs of cells interacting with parameters from the points indicated as 1 to 4 in (A). Parameter values in point 1 correspond to those used in Collier et al. (1996) (B), while parameter values in points 2-4 (C-E) correspond to examples of other asymmetric pairs, and symmetric positive and symmetric negative pairs.

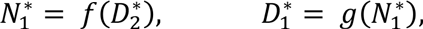

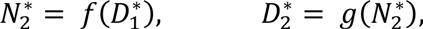

In parallel with this classification of steady states, a cell is considered to be Notch positive when the level of Notch surpasses a certain threshold *N_thr_* (considered here to be 0.1), and Notch negative in the opposite case. In the case of the *Drosophila* midgut, a Notch positive cell would correspond to an EB, and a Notch negative cell to an ISC. In that way, a homogeneous steady state can represent either an ISC/ISC pair (symmetric Notch negative) or an EB/EB pair (symmetric Notch positive). Most heterogeneous steady states, in turn, correspond to an ISC/EB pair, although heterogeneous states in which both values of Notch lie below (or above) the threshold *N_thr_* still represent symmetric ISC/ISC (or EB/EB) pairs. This is reflected in Fig. 2A through the difference between the stability boundary (dotted grey line) and the boundaries between the fate-pair domains shown in colour code.

### Dynamical behaviour

We investigate the temporal evolution of the model by solving numerically the Equations (6). For this purpose we use a finite difference approximation (two-stage Runge-Kutta; LeVeque, 2007). In our calculations, we are considering *r=h=2* and *δ_N_=δ_D_*, as in previous works (Collier et al., 1996; Sprinzak et al., 2010). Cells are considered initially as negative for Notch activation, with similar initial levels of Notch and Delta (*N_1_(t=0)=0.05, N_2_(t=0)=.06, D_1_(t=0)=0.90, D_2_(t=0)=0.91* in dimensionless units).

### Drosophila culture and strains

Adult flies were raised in standard cornmeal medium, collected daily and maintained in fresh vials with added yeast (food replaced every 24-48h) until dissection at 4-6 days of age.

EBs were identified by co-expression of an enhancer trap reporter of the undifferentiation marker *escargot (* Fly Base: *P{PTT-GB}esg^YB0232^;* Quiñones-Coello et al., 2007) and the synthetic Notch transcriptional activity reporter *GBE-Su(H)-LacZ* (FlyBase: *P{Ddc.E(spl)m8-HLH-lacZ.Gbe};* Bray and Furriols, 2001), while ISCs were identified by expression of the *esg* reporter alone (Micchelli and Perrimon, 2006; Ohlstein and Spradling, 2006) (Fig. 1A).

### Immunohistofluorescence and imaging

Immunofluorescence was performed essentially as described in (Bardin et al., 2010) but with a heat-fixation step (Miller et al., 1989).

Primary antibodies were: chicken anti-*β*-Galactosidase (Abcam ab9361, 1:200), rabbit anti-GFP (Abcam ab6556, 1:200), anti-Arm (mAb N2-7A1, Developmental Studies Hybridoma Bank, 1:50), sheep anti-Notch (Muñoz-Descalzo et al., 2011; 1:1000). Secondary antibodies conjugated with Alexa fluorophores were from Invitrogen (1:500).

Confocal stacks were obtained in a Zeiss LSM 710 with an EC Plan-Neofluar 40X oil immersion objective (numerical aperture 1.3), with voxel size 0.21x0.21x1.00 or 0.14x0.14x0.42 µm (XYZ) for the quantification of contact area (with no oversampling in Z) and Notch distribution, respectively.

### Image analysis

To measure contact area, stacks were analysed with a combination of ImageJ macros and python scripts to (1) manually identify all esg-GFP^+^ cells in a z-projection of the stack, (2) automatically threshold *GBE-Su(H)-lacZ* reporter expression in 3D to determine its expression status (positive or negative) in every esg^+^ cell, (3) manually identify the nests of esg^+^ cells so that (4) a series of 3D stacks, containing only one pair each, is automatically cropped, and (5) the contact membrane of each esg^+^ cell pair is semi-automatically determined using FIJI for each optical plane, by binarising the immunofluorescence of Armadillo/*β*-catenin (Arm). Arm labels the membrane throughout the apical-basal axis (see Results), which allows measuring the amount of contacting membrane in each cell pair as the number of Arm^+^ voxels shared between the two cells (expressed in *µ*m^2^).

For measuring Notch and Arm distribution at the membrane, the membrane contours (3 pixels wide) of cells in pairs were manually determined in each plane. Intensity data from those positions were used as follows:

*Intensity normalisation*. For each cell pair, two nearby 50x50 pixel squares spanning the full z-stack were manually selected, the signal therein averaged for all the planes where membrane was detectable, and this average value taken as background. Notch and Arm intensity values for that stack were normalised by dividing by the background value.

*Distribution along cell perimeter*. Each membrane pixel position was assigned an angular value respect to the centroid by calculating its tangent arc (±π depending on the quadrant). Thirty overlapping sliding windows (of 2π/15 rad with half window overlap) were delimited in each plane, and their pixel intensities were normalised and averaged.

*Distribution along the apical-basal axis*. Each cell was sliced in 10 overlapping angular windows (2π/5 rad with half overlap). For each window, a normalised, average intensity measurement was taken per confocal plane (i.e. along the apical-basal axis). Apical-basal positions were normalised from 0 to 1. Intensity data points along the apical-basal axis were obtained by interpolation from average normalized intensity values.

## Results

### Lateral inhibition can result in stable, opposing symmetric signalling states

We study the steady-state behaviour of a standard model of lateral inhibition for the case of two cells (see Methods). The steady states of this system depend on two parameters, *a* and *b* (the dimensionless activation and inhibition thresholds, respectively; see Methods), which we allow to vary across the population of cell pairs. We then calculate the equilibrium state of the system in this two-dimensional parameter space, according to the resulting signalling profile: asymmetric (one cell positive for Notch activation and the other one negative, see Methods), symmetric positive, or symmetric negative for Notch activation (Fig. 1E). Thus, for a population of cell pairs with variable activation or inhibition thresholds (*a* and *b*), the three possible signalling state profiles occur (Fig. 2A, see Fig. 2B-E for a comparison of the dynamic evolution of examples of the three profiles, with Fig. 2B corresponding to the parameter values from Collier et al., 1996). The three signalling state profiles can be found within a relatively short range of parameter values (Fig. 2A), and this scenario does not change qualitatively when considering a wide range of threshold values for Notch activity classification, as defined in the Methods section *(0.001 ≤ N_thr_ ≤ 0.7)* (Fig. S1).

In a biological system, the existence of three possible signalling state profiles would be equivalent to having three different cell fate combinations across a population of initially uncommitted cell pairs interacting through Notch/Delta, with the specific fate combination of a given cell pair depending on the sensitivity to Delta activation and Notch inhibition of the pair. To investigate the potential of this lateral inhibition model, incorporating variable activation and inhibition thresholds, to describe a real biological system, we turned to the *Drosophila* midgut.

### Cell contact area as indicator of activation threshold

In the *Drosophila* midgut, Notch negative cells correspond to ISCs, and Notch positive to EBs. The Notch activity reporter *GBE-Su(H)* is hardly expressed above background levels in ISCs (Ohlstein and Spradling, 2007), and our own observations), and hence our choice of a low threshold value, *N_thr_ = 0.1*. Symmetric positive pairs will equate to an event of symmetric differentiation (EB/EB), symmetric negative pairs to symmetric self-renewal (ISC/ISC) and asymmetric pairs to asymmetric ISC fate (ISC/EB) (Fig. 2A).

To relate the model to real tissue, we need first to consider how the dimensionless parameters *a* and *b* are related to biological features displaying variability across undifferentiated (esg^+^) cell pairs. We assume that biochemical processes intrinsic to the cell, such as protein degradation rates (*δ_D_* and *δ_N_*), the maximal biosynthesis rates (α and β), and the threshold of Delta inhibition by Notch in the same cell (*K_D_*), will not be highly variable among cells with a common developmental identity. On the other hand, the threshold of Notch activation by neighbouring Delta (*K_N_*) depends directly on the interaction between the two cells, which could be variable for different pairs of cells. For instance, due to the spatial constraints of cell packing, the contact area between cell pairs could be substantially different from pair to pair. Indeed, tissue images reveal that undifferentiated cells in nests show irregular shapes and variable contact area (Fig. 3A). In the Notch/Delta system the amounts of Notch and Delta are usually limiting, and this seems to hold true for the adult *Drosophila* gut, where haploinsufficiency has been described (Biteau et al., 2008; de Navascués et al., 2012). Therefore, it is expected that variations of ∼2-fold or more in contact area would lead to significant changes in the levels of Notch activation. This is captured in the model by the dimensionless activation threshold *a*, which can be assumed in a first approximation, following Eqn 7 above, to be inversely related with the contact area (the larger the contact area, the easier it is for Delta to activate Notch, and thus the smaller the activation threshold). As shown in Fig. 2A (see also Fig. S2), variation in *a* best allows for heterogeneity in stable steady state levels of Notch activity and therefore in fate choice. From this we hypothesize that variation of contact area (or any other biological feature correlating with the threshold of Notch activation) is likely to allow the diversity in fate outcome that we observe in the model.

For this relationship to hold, Notch distribution at the cell membrane must be such that the amount of receptor correlates with the contact area. To evaluate this, we examined the localisation of both Notch and Armadillo/*β*-catenin (Arm) in both single and paired undifferentiated cells (recognised by *Notch* expression; Bardin et al., 2010) of the adult posterior midgut epithelium with confocal microscopy (Fig. 3A). In *Drosophila* epithelia, Arm/*β*-cat participates in the formation of adherens junctions, which localise mostly in an apical domain, with lower levels at the lateral membrane (Tepass and Hartenstein, 1994). Therefore, we used Arm staining to define membrane boundaries in 3D, and measured the intensity of Notch and Arm signals at the cell membrane.

We could not find any strong pattern in the variations of Notch immunodetection intensity within confocal planes, and it would only seem that Notch is slightly enriched at the boundary between two esg^+^ cells (Fig. 3B). This indicates that Notch concentration is largely independent of the position at the membrane along the cell perimeter, and in particular along the contact between esg^+^ cells. Moreover, the localisation of Notch along the apical-basal axis of the cells is also largely homogeneous. This is manifest in the small variation in the average amounts of Notch between different optical planes (Fig. 3D), and in the narrow distribution of mean values per plane, with low values of coefficients of variation per plane, of Notch intensity values (Fig. S3B,D). Therefore, the contact area between cells is a good approximation to the total amount of Notch receptor available for signalling.

We note that Arm largely parallels Notch localisation at the membrane (Figs. 3C,E and S3A,C) but shows a stronger enrichment at the boundary (Fig. 3C), in agreement with previous reports (Maeda et al., 2008). Incidentally, these results also reveal that neither Arm nor Notch are restricted to the apical domain in the midgut epithelium, and instead can be found in similar amounts along the apical-basal axis of the membrane in ISCs and EBs (Fig. 3D-F). This situation contrasts with Arm and Notch distribution in other *Drosophila* epithelia (Tepass and Hartenstein, 1994; Tepass et al., 2001).

**Figure 3.**
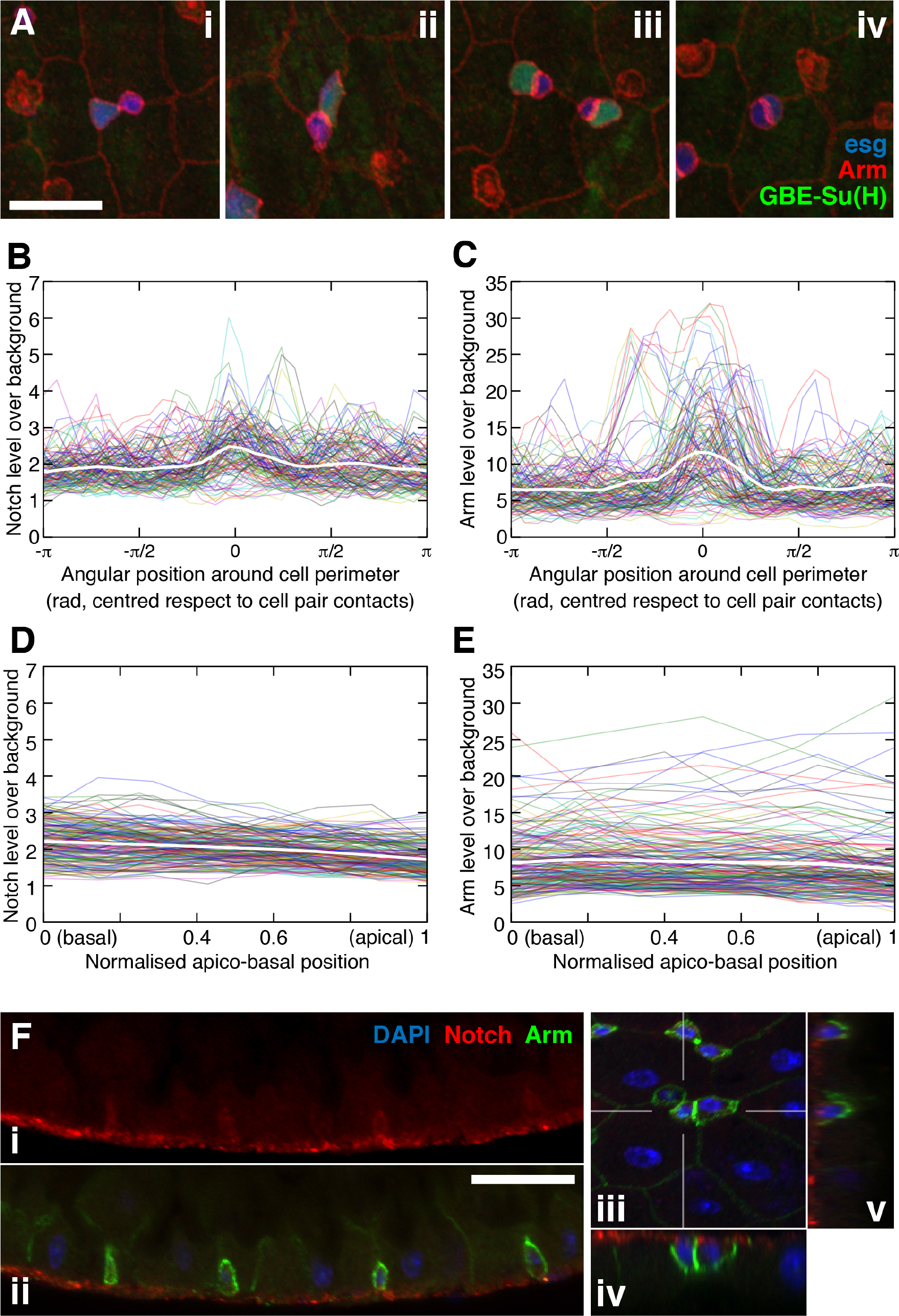
Variability in contact area and distribution of Notch at the membrane. A. Confocal stacks projected in Z, showing variability in contact length (as proxy for area). Scale bar: 20*µ*m. B-C. Notch (B) and Arm (C) levels along the perimeter of the cell planes (colour lines) and mean (white). For each cell plane, position 0 corresponds to the centre of the contacting membranes (defined as the position that intersects the line connecting the cell centroids in that plane). D-E. Notch (D) and Arm (E) levels along the apical-basal cell axis (with height of the cell normalised to 1). Each cell contributes ten lines to the plot, corresponding to the intensity values along the vertical axis of non-overlapping, angular windows of 2π/10. Data displayed in B-E are from 20 paired esg+ cells. Data in B-E are from 20 paired cells. F. (i, ii) Side views of the intestinal epithelium, showing the apical-basal distribution of Notch and Arm. Lumen is at the top and basal at the bottom. (iii-v) Top view of the intestinal epithelium (iii) with ZY and XZ side views (iv, v) corresponding to the marked lines in (iii). Scale bar in (ii): 10*µ*m.

Taken together, our results suggest that Notch receptor is randomly distributed in the cell membrane, which suggests that measurements of membrane contact area may be relevant to the dynamics of Delta-Notch signalling as a proxy for the activation threshold *a* in our model.

### Correlation of contact area values and cell fate profiles

We have shown that contact area can be used as a measure of the amount of Notch available for interaction with Delta. We thus measured contact area in 508 pairs of esg^+^ cells with both symmetric (ISC-ISC and EB-EB) and asymmetric (ISC-EB) fates (Fig. 1D-E); these pair classes have been described in (de Navascués et al., 2012; Goulas et al., 2012). We found the contact area in these pairs to be highly variable, ranging from just around 1*µ*m^2^ to over 60*µ*m^2^ (Fig. 4A) and with a high coefficient of variation (0.52). This degree of variability indicates that contact area has the potential to be a regulatory mechanism of the system (through its influence on the dimensionless activation threshold *a*).

**Figure 4.**
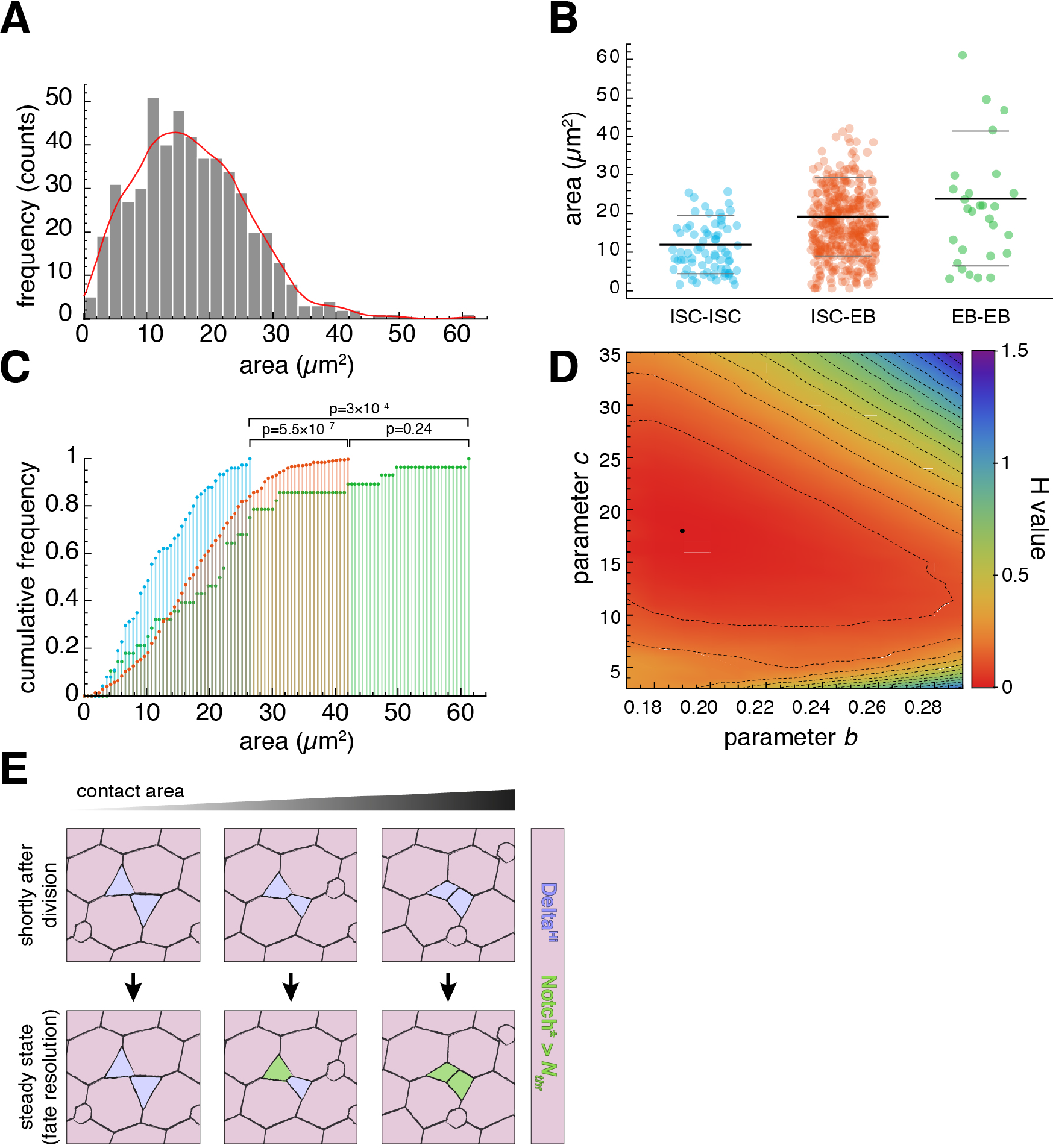
The model can reproduce the observed cell fate profiles. Data set from N=508 nests (Fig. 1D). A. Frequency of contact area values for nests of two undifferentiated cells. Line marks the Smooth Kernel Distribution (SKD) used to generate areas for the simulation. B. Contact area values segregated by fate profiles. Black lines indicate mean values. Grey lines mark the position of one standard deviation below or above the mean. C. Cumulative frequency of the contact area data displayed in (B). Note that ISC-EB and EB-EB distributions cannot be distinguished statistically. D. Kullback-Leibler relative entropy (H) between experimental and model distributions as a function of b and c. Values of area in the model are generated by the SKD depicted in (A). Best value corresponds to b=0.19 and c=18 (black dot) and leads to fate profile proportions as in Table 1. E. Fate outputs for the lateral inhibition model for three different values of the contact area.

Our model predicts that a biological feature influencing Notch/Delta signalling thresholds in cell pairs should correlate with the patterns of symmetric and asymmetric fates. Therefore, we classified measurements of contact area according to the fate profile of their corresponding cell pair and compared their values (Fig. 4B). We found that on average, the contact area between ISC-ISC (11.59 ± 0.73 *µ*m^2^; mean ± standard error of the mean) is clearly smaller than those of ISC-EB (17.68 ± 0.42 *µ*m^2^) and EB-EB pairs (21.6 ± 2.76 *µ*m^2^). This is also clear when considering the distribution of sizes for each pair type and confirmed by the Kolmogorov-Smirnov test (Fig. 4C). From these results we take that ISC-ISC pairs have a significantly smaller contact area than the other two fate profiles.

### Updating the model with area variation reproduces fate profile distributions

This finding gives us biological justification to consider the activation threshold *a* to be variable in the model (inversely proportional to the contact area), and test the capacity of the model to produce the observed proportions of fate pairs. To do this, we first generated a large sample of contact area values *A*, from a Smooth Kernel Distribution based on the experimental data (Fig. 4A). To input values from *A* into the model, we considered the area values *A* inversely related to the activation threshold *a* by a constant *c* (*a* = *c*/*A*), treated as a parameter of the model. We then analysed the stable steady states of the model, obtaining the proportions of the three possible fate pairs resulting from *A*, for different values of *b* and *c*.

In order to compare the fate distribution obtained from the model and the experimental data we use the Kullback-Leibler relative entropy (*H*), which is a dissimilarity measure between two probability distributions (giving the value 0 if the distributions are equal; Kullback and Leibler, 1951). We found an excellent agreement between the proportions of EB-EB, EB-ISC and ISC-ISC pairs observed experimentally, and the distributions from the model for an extended range of values of *b* and *c*, as indicated by the low values of *H* between theoretical and experimental distributions (Fig. 4D). The best fit (Table 1) is obtained with *b*=0.19 and *c*=18 (*H*=0.3×10^-3^, black dot in Fig. 4D). By mapping *A* input values (Fig. 4A) for *b*=0.19 and *c*=18 to the model phase diagram (horizontal line in Fig. 2A), one finds that ISC-ISC pairs occur at the lowest values of contact area, in good agreement with our experimental observations (Fig. 4B-C). In this region of the parameter space, EB-EB and ISC-EB pairs are found at intermediate or high values of contact area (lower *a*), respectively. While the model favours EB-EB pairs resulting from smaller contact areas than ISC-EB pairs, we cannot distinguish statistically between the two experimental distributions of contact area (see Fig. 4C). Hence, we propose that the contact area between pairs of cells can influence the fate outcome of Notch/Delta signalling in the *Drosophila* midgut (Fig. 4E), with small contact area clearly favouring symmetric self-renewal.

**Table 1.** Cell-fate profiles as obtained experimentally and theoretically. The value of Kullback-Leibler relative entropy between the two distributions is 0.3 x 10^-3^. Model parameters: *b* = 0.19 and *c* = 18.

| **pair fate** | **experimental** | **model** |
| --- | --- | --- |
| ISC-ISC | 14.6 % | 15.1% |
| EB-EB | 5.5 % | 5.0 % |
| ISC-EB | 79.9 % | 79.9% |

## Discussion

We have considered a standard model of Notch/Delta-mediated lateral inhibition (Collier et al., 1996) and investigated the effect of the trans-activation of Notch by Delta and the inhibition threshold of Delta by Notch signalling (here considered phenomenologically as the dimensionless thresholds *a* and *b*, respectively) on the dynamics of lateral inhibition for a system of two cells. We find that, provided there is a degree of variability in signalling thresholds between cell pairs, three different signalling states (and therefore fate combinations) can occur under the same conditions. This is a considerable expansion of the model, whose use has so far been mostly centred on solutions that provide fine-grained (checkerboard) patterns. This population-level variability of signalling thresholds can be associated to diversity in the contact areas between pairs of cells. The model reproduces the signalling outcomes observed in the *Drosophila* intestine, which translate into differentiation vs. self-renewal fates. This in turn provides a mechanism whereby ISCs may undergo neutral competition, which is a widespread pattern of adult tissue maintenance in metazoans from *Drosophila* to humans. These results thus provide a potential biological justification for the neutral competition of *Drosophila* adult ISCs.

The seminal work by Collier et al. (1996) established a minimal model which crystallised the biological intuition of lateral inhibition developed from the observation of neurogenesis and akin processes of precursor selection (Sprinzak et al., 2011; Formosa-Jordan et al., 2012; Petrovic et al., 2014): amplification of small differences in signal, leading to checkerboard patterns of stable, all-or-none signalling states. This formalisation, with their parameter choice for *a*, *b*, has justly become a reference in the field. In the last few years there have been expansions of this model to accommodate additional features, such as the specific activity of Jagged, another Notch ligand (Boareto et al., 2015), or departures from fine-grained patterns based on different ratios of positive and negative Notch cells (de Back et al., 2013). These models introduce additional genetic components or noise terms that allow the new phenomenology. By contrast, we have left intact the general dynamics of the minimal model of Collier et al. (1996) and introduced only a degree of variability in the sensitivity of each cell pair to signal transduction.

Another important contribution of Collier et al. (1996) was to identify the mathematical condition for symmetry breaking. We explore this condition further and determine systematically the contribution of the signalling thresholds *a* and *b* to the condition of stability (Fig. 2A). It would also be interesting to explore how variation in the cooperativity of the Notch trans-activation or Delta inhibition (parameters *r*, *h*) affect the capability of the system to arrive to symmetric or asymmetric steady states, as seen recently in Turing patterns (Diambra et al., 2015).

Our work considers the contact area between cells engaged in signalling as the source of variation in signalling threshold. Contact area can be an effective tuning parameter of a biological system (Khait et al., 2015), since it can integrate mechanical constraints into signalling, as it has been shown for cell density and proliferative control by the Hippo pathway (Schlegelmilch et al., 2011; Kim et al., 2011; Silvis et al., 2011). In a system such as the posterior midgut, where some differentiated cells are much larger than their progenitors (see Fig. 1A), differentiated and mature cell loss certainly would have a local impact in the packing geometry of cells interacting via Notch/Delta, and connect naturally with the fate outcome of stem cell divisions. This could be particularly useful in conditions of regeneration. Importantly, our theoretical framework could in principle accommodate any source of variation; for instance, variation arising from the unequal (either random or regulated) inheritance of signalling components could result in variation in the capability of signal transduction in the population. It is interesting to consider that while shortly after division most of the ISC daughter cells display similar levels of Notch and Delta proteins (Ohlstein and Spradling, 2007), endosomes bearing the signalling molecule Sara display an inhomogeneous inheritance pattern (Montagne and González-Gaitán, 2014). It has recently been found that ISC divisions producing enteroendocrine cell precursors do segregate Delta asymmetrically towards the precursor cell (Guo and Ohlstein, 2015), which suggests that ISCs switch between different types of cell division.

Understanding how Notch/Delta signalling results in stochastic cell fate patterns is of particular relevance in adult homeostatic tissues, as Notch signalling controls fate in many types of tissue stem cells (Koch et al., 2013). Moreover, many adult stem cells balance their fate via neutral competition (Krieger and Simons, 2015). Our model proposes a mechanism whereby Notch/Delta signalling could result in neutral competition of stem cells by lateral inhibition between sibling cells. This provides an alternative explanation to the neutral competition of Drosophila adult ISCs, which has been proposed to arise from Notch/Delta-mediated lateral inhibition involving the offspring of non-related ISCs, coinciding in space (de Navascués et al., 2012) and resolving 20% of the time in symmetric fate. However, that proposal faces the difficulty that ISC/EB nests rarely contain more than two cells (de Navascués et al., 2012). Moreover, we and others have found isolated pairs of ISCs or EBs frequently in the tissue (de Navascués et al., 2012; Goulas et al., 2012). Our model provides a potential explanation of how the offspring of a single ISC (pairs of Notch/Delta signalling cells) may reach a symmetric steady state, leading to symmetric self-renewal o differentiation.

It would be interesting to see how our model (based on signalling threshold variability) translates to a larger group of interacting cells, in particular in light of recent findings in the esophageal epithelium. There, tissue is maintained by the neutral competition of basal progenitor cells (Doupé et al., 2012), and this competition is heavily influenced by Notch signalling, to the point that alterations in the pathway can lead to the fixation of mutant clones and poise the tissue for tumour initiation (Alcolea et al., 2014).

## Acknowledgments

We wish to acknowledge support from CONICET to NG and from Cardiff University to JdN. JGO and RMC are supported by the Spanish Ministerio de Economía y Competitividad and FEDER, through project FIS2012-13360-C02-01, and the ICREA Academia Programme. RMC also acknowledges financial support from La Caixa Foundation. We would like to thank Alfonso Martínez Arias for his encouragement and support and Nicole Gorfinkiel and David Sprinzak for critical comments on the manuscript.

**Competing Interests**

No competing interests declared.

---

**Supplementary Figure 1.**
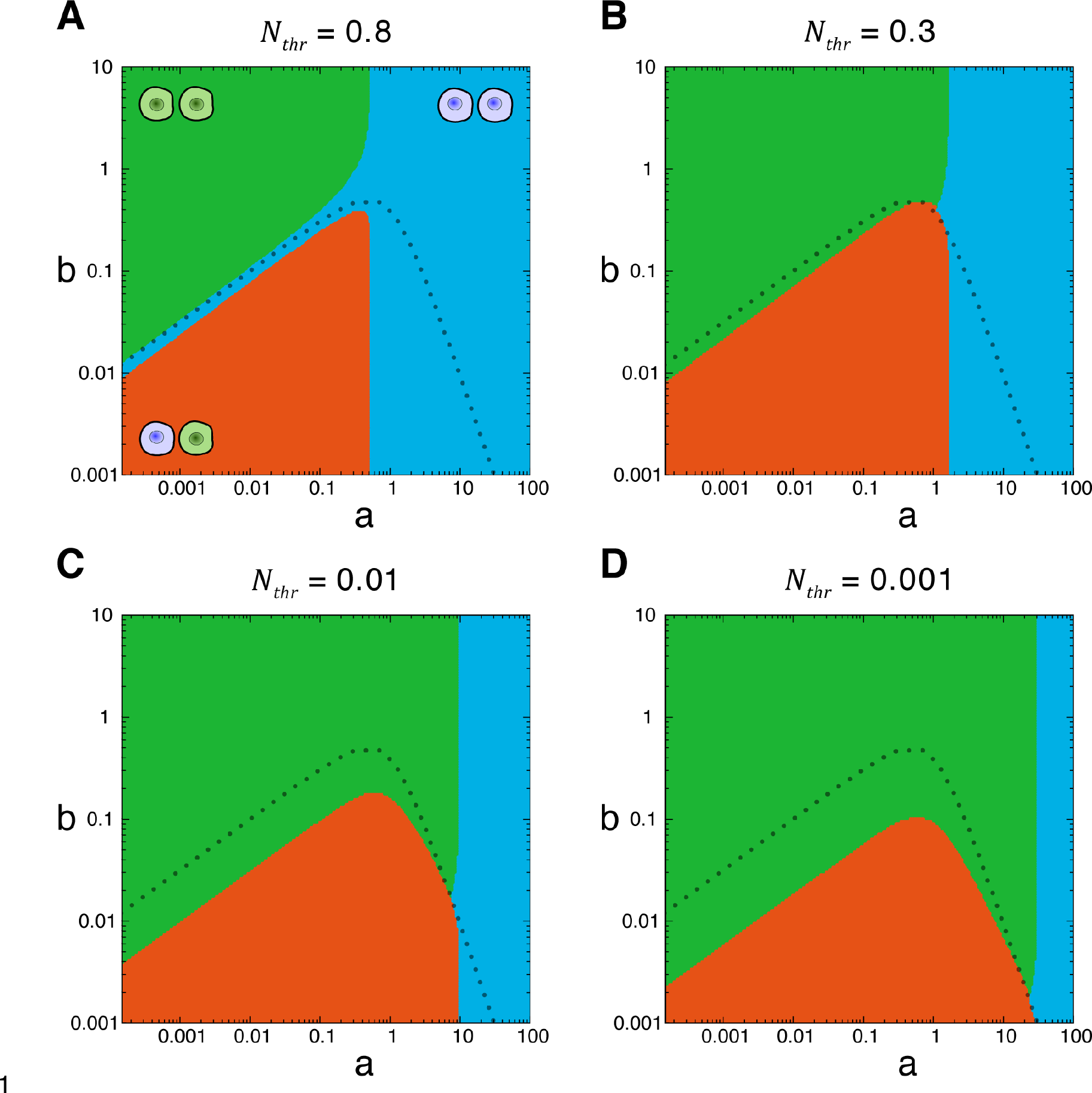
Fate profiles in parameter space over a broad range of threshold *N_thr_* values. **A-D**. Phase space for *N_thr_* equal to 0.8 (A), 0.3 (B), 0.01 (C), and 0.001 (D), respectively. The dotted line marks the stability boundary for the ‘homogeneous’ solutions (pairs of identical cells), and serves as reference for comparison with Figure 2A. While in A (where *N_thr_* > 0.7), the area of asymmetric fate is surrounded by symmetric negative resolution, in B-D the organisation of the phase space is very similar, with the transitions shifting along the stability boundary.

**Supplementary Figure 2.**
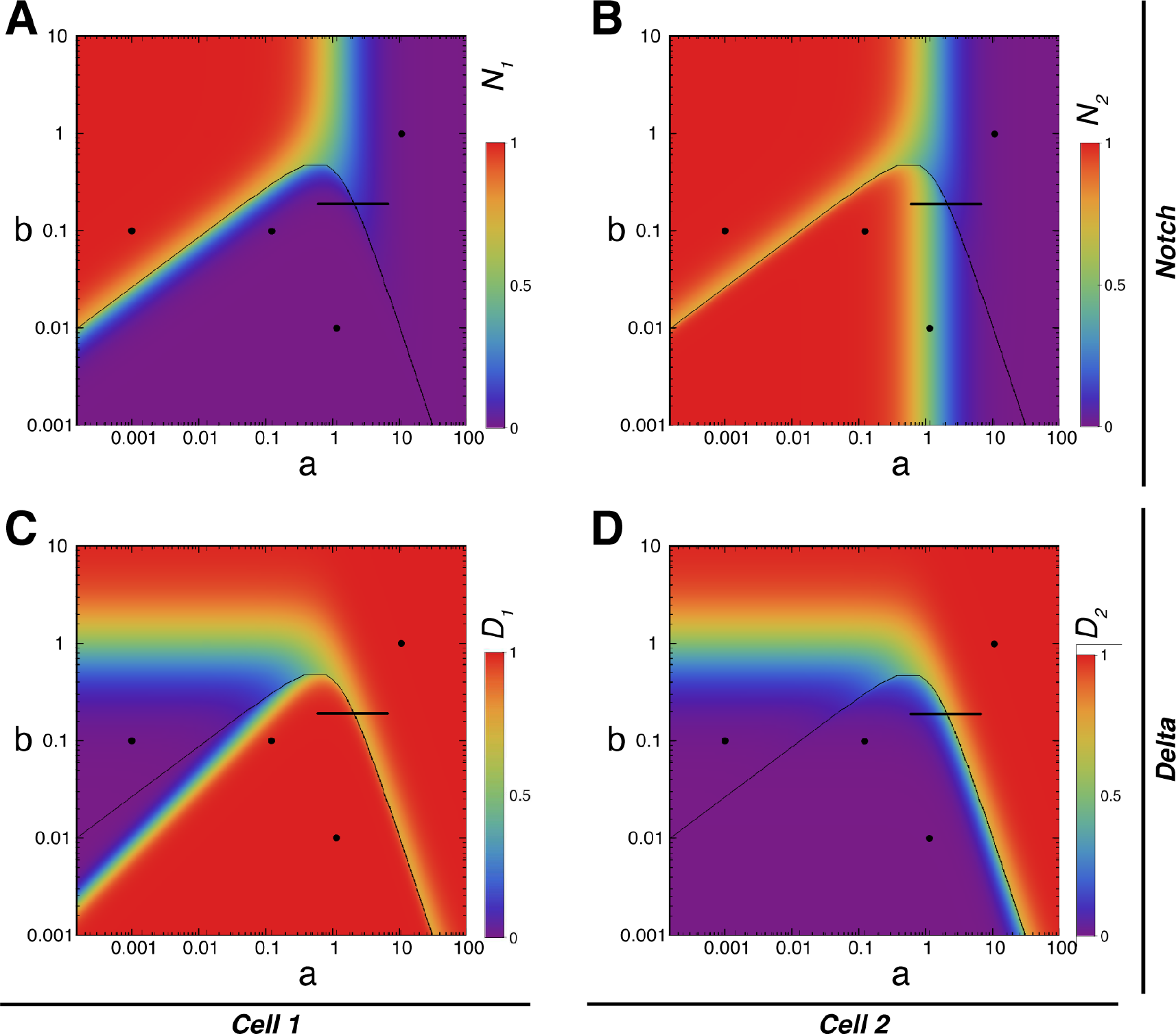
Values of Notch and Delta at steady state across parameter space. The short, horizontal line indicates 95% of “a” values used to generate a theoretical distribution of cell fates for b=0.19 (see main text). Dotted line, boundary of stability for steady states with identical cells. The black dots mark the parameter values used in (Collier et al., 1996) (Figure 2B) and the asymmetric, symmetric positive and symmetric negative pairs from Figure 2C-E. **A, B**. Steady-state values of activated Notch the two cells of a pair (one in each panel) respect to *a, b*. **C-D**. Steady-state values of Delta in the two cells of a pair (one in each panel) respect to *a, b*. Note that depending on the value of activated Notch, one can find symmetric negative or symmetric positive fate profiles below the boundary (region of heterogeneous solution), showing that the model allows for symmetric steady states where cells in a pair do not have identical amounts of Notch or Delta.

**Supplementary Figure 3.**
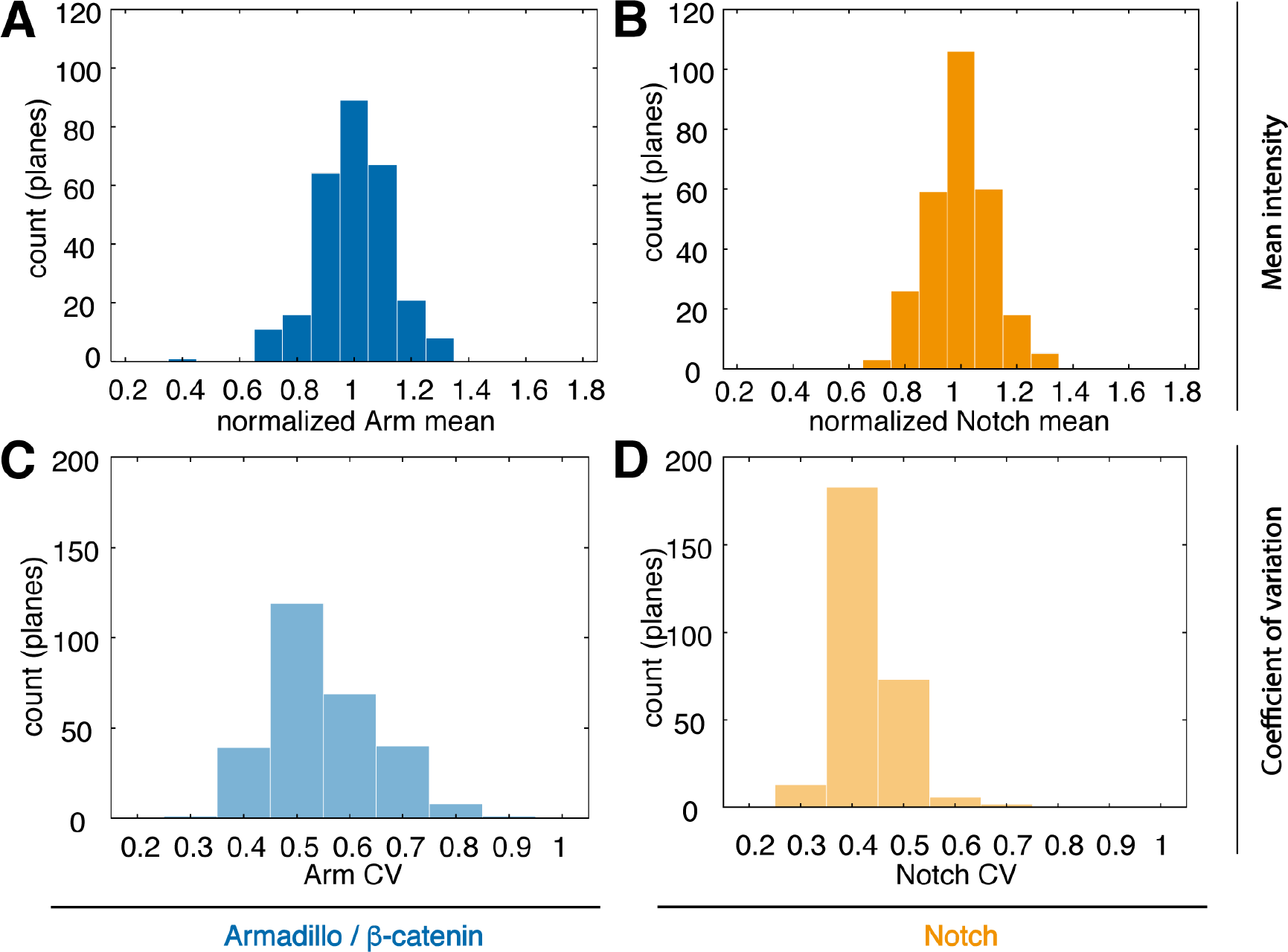
Distribution of Notch and Armadillo at the membrane. Data correspond to all 46 analysed cells (single and paired). **A-D**. Histograms of the normalised mean intensity per plane (A, B) and the coefficient of variation (CV) per plane (C, D) for Notch (A, C) and Armadillo (B, D) markers. The normalised mean intensity in plane *i* is defined as the ratio of the average of the plane and the average for the cell.

